# Neuroimaging and plasma biomarker differences and commonalities in Lewy body dementia subtypes

**DOI:** 10.1101/2025.01.24.634687

**Authors:** Naomi Hannaway, Angeliki Zarkali, Rohan Bhome, Ivelina Dobreva, George EC Thomas, Elena Veleva, Irene Gorostiaga Belio, Katie Tucker, Amanda Heslegrave, Henrik Zetterberg, Rimona S Weil

## Abstract

**INTRODUCTION:** Despite ongoing debate about whether Parkinson’s disease dementia (PDD) and dementia with Lewy bodies (DLB) are separable diseases or a single Lewy body dementia (LBD) spectrum, there are limited neuroimaging investigations of differences between these conditions.

**METHODS:** We used fixel-based diffusion MRI and plasma measures to examine white matter integrity and burden of amyloid pathology (using tau phosphorylated at theonine-217 (p-tau217) in 47 patients with DLB, 21 PDD, 29 PD and 23 age-matched controls.

**RESULTS:** We show reduced fibre cross-section in LBD versus PD, and increased concentrations of plasma neurofilament light chain and p-tau217; with p-tau217 and fibre cross-section associated with cognition. Fibre density was reduced in PDD versus DLB, but neither plasma measures nor fibre cross-section differed between LBD subtypes.

**DISCUSSION:** Our findings suggest differences in white matter integrity between DLB and PDD that are driven by distinct processes from those causing changes in white matter integrity in LBD compared with PD.

## 1. Background

Lewy Body Dementia (LBD), comprising Parkinson’s disease dementia (PDD) and Dementia with Lewy bodies (DLB), is the second commonest degenerative cause of dementia after Alzheimer’s disease.^1^ PDD and DLB are characterised by overlapping symptoms of dementia, motor parkinsonism, visual hallucinations, cognitive fluctuations and rapid eye movement (REM) sleep behaviour disorder.^2^ They also share overlapping pathologies, including cortical build-up of alpha-synuclein, beta-amyloid plaques and tau tangles.^3,4^ Currently, differences between DLB and PDD are defined clinically, based on the ‘one year rule’. If dementia precedes or develops within one year from onset of motor symptoms, DLB is diagnosed, whilst if dementia occurs more than a year after onset of motor Parkinsonism, PDD is diagnosed. There is continued debate whether DLB and PDD are part of a single disease spectrum,^5,6^ or whether these are two separable diseases.^7^ However, few studies directly compare these two patient groups.

Fluid biomarkers and advanced brain imaging techniques provide complementary information, which together offer potential to shed light on disease-related changes in LBD. Diffusion weighted imaging (DWI) is sensitive to changes in white matter connections, reflecting axonal damage that is observed before grey matter atrophy^8^ Measured using DWI, fractional anisotropy (FA) is reduced in LBD relative to age-matched controls in regions including corpus callosum, superior longitudinal fasciculus and parietal lobes.^9–12^ Comparisons between LBD subtypes are less clear. Whilst some studies found no differences,^11^ others observed widespread changes in DLB compared to PDD, with reduced FA in visual association, posterior temporal and posterior cingulate fibres.^12^

However, conventional diffusion imaging has limited sensitivity in regions with crossing fibres, which are present in the majority of the brain.^13^ Higher-order diffusion models, such as fixel-based analysis, overcome this by examining individual fibre populations (fixels) within a voxel,^14^ increasing sensitivity in regions with crossing fibres. We previously showed in PD that those at high-risk for dementia have widespread white matter changes compared with those at low-risk for dementia, with greater loss of fibre cross-section associated with poorer cognition after 18-months^15^. Whether these differences are also seen between LBD subtypes is not yet established.

Widening availability of fluid markers enables complementary information about pathological accumulations and other markers of disease to be collected. One such protein, neurofilament light chain (NfL) is a marker of neuronal injury^16^, reflecting axonal damage^17^. Increased plasma NfL is seen in people with DLB relative to age-matched controls^18^ and PDD compared to PD^19^, with higher concentrations associated with poorer cognition^19^ and white-matter loss^8^. Co-pathology is frequently seen in LBD with beta-amyloid and tau the commonest pathological accumulations^3^. These can be detected with plasma tau phosphorylated at threonine 181 (p-tau181) and at threonine 217 (p-tau217), which are both highly sensitive to brain beta-amyloid and tau, and established as markers of beta-amyloid and tau pathology^20,21^. Plasma p-tau181 is increased in DLB compared with controls^18,22^, and associated and with poorer cognition^22^. In PDD, results are less consistent, with no differences in PD between patients with and without dementia^19^. However, p-tau217 has not yet been tested in PDD.

Establishing whether DLB and PDD differ in white matter connections, and how these relate to plasma markers has potential clinical implications as it could determine whether particular patients would benefit from a common approach or from more targeted interventions.

Here, we used fixel-based analysis on diffusion-weighted MRI; and plasma NfL and p-tau217, as markers of axonal damage and beta-amyloid and tau co-pathology, in people with DLB and PDD and PD with normal cognition who were also characterised as low-risk for developing PD dementia. We predicted greater loss of white matter integrity and increased concentrations of NfL and p-tau217 in people with LBD compared to PD and age-matched controls. Based on previous work using less sensitive FA measures, we predicted greater loss of white matter integrity and higher NfL in DLB than PDD^12^; and based on reports of increased beta-amyloid pathology^23^ in DLB, we expected higher levels of p-tau217 in DLB than in PDD.

## 2. Materials and methods

### 2.1 Participants

People with DLB, PDD, PD and age-matched controls aged 50–81 years were recruited from the National Hospital for Neurology and Neurosurgery outpatient clinics and affiliated hospitals, from national patient support groups (Lewy Body Society and Rare Dementia Support), and from an existing observational cohort study of PD (led by RSW) (see e.g. Hannaway 2023^24^ for more details).

DLB and PDD participants had a clinical diagnosis according to Diamond Lewy toolkits for DLB or PDD ^25^. People with PD had a clinical diagnosis of PD according to the Movement Disorders Society (MDS) clinical diagnostic criteria.^26^ We further categorized people with PD into high and low-risk for dementia, based on their performance on two computerised visual tasks: biological motion^27^ and the “Cats and Dogs task”^28^. We have previously shown that poor performance on these tasks predicts dementia and poor outcomes in PD^15,24^; and that people with PD who are high-risk for dementia already show extensive loss of white matter integrity, and changes in plasma markers^8,15^. We therefore excluded people with PD categorised as high-risk for dementia, based on performance in these tasks.

All patients were within 10 years of their respective diagnosis. For PDD, this was within 10 years of the dementia diagnosis. Where people with PD or controls had attended for more than one visit in the longitudinal study, the last completed visit without dementia or mild cognitive impairment (MCI) was used. This was done to match ages to people with LBD. Where people with PD in the longitudinal study had developed PD-MCI or dementia, their latest visit was used, and they were counted as PDD.

People with a history of confounding neurological or psychiatric disorders were excluded, as were metal implants considered unsafe for MRI. Controls diagnosed with dementia or mild cognitive impairment, or with a Mini Mental State Examination (MMSE) score of less than 25 were also excluded.

All participants gave written informed consent, and the study was approved by the Queen Square Research Ethics Committee (15.LO.0476).

### 2.2 Clinical and neuropsychological assessments

Participants underwent detailed clinical and neuropsychological assessments. The Mini-Mental State Examination (MMSE) and Montreal Cognitive Assessment (MoCA) were completed as measures of global cognition, plus 2 tests per cognitive domain. These were: Attention: Stroop colour naming^29^ and digit span^30^; Executive functions: category fluency^31^ and Stroop interference^29^ Language: letter fluency and graded naming task^32^; Memory: word recognition task^33^ and logical memory^30^; Visuospatial function: Hooper visual organization test^34^ and Benton Judgement of line orientation.^35^

Due to higher frailty in people with LBD, it was not always possible to complete the full battery of neuropsychological testing during the study visit. Where this was the case, the MMSE and MoCA were prioritized, followed by one task per cognitive domain (in each case, the first of the two tests listed above was used). Furthermore, if participants were unable to complete the full Stroop task, they completed a “Half-Stroop” consisting of the first 3 lines of the task.

A summary cognitive score was calculated as the averaged z-scores of the MoCA plus one task per cognitive domain: namely inverted Stroop (colour naming time), category fluency, letter fluency, Recognition Memory Test and Hooper Visual Organization Test, as we have previously described^24^. Where only Half-Stroop was completed, time to complete the full-Stroop was predicted using a regression model to allow calculation of the summary cognitive score (further details in Supplementary Methods, Supplementary Figure 1).

Motor symptom severity was measured using the United Parkinson’s Disease Rating Scale, part 3 (UPDRS-III)^36^, and the timed up and go test (TUG)^37^. Total symptoms were measured using the UPDRS parts I, II, III, and IV. For all clinical and cognitive measures, participants were tested whilst taking their usual medications, and Levodopa Equivalent Daily Dose (LEDD) was calculated^38^.

Cognitive fluctuations were measured using the clinician assessment of fluctuations (CAF), one-day fluctuations scale^39^ and the dementia cognitive fluctuation scale (DCFS)^40^. Anxiety and depression were measured using the Hospital Anxiety and Depression Scale (HADS)^41^. Impairments in activities of daily living were measured using the Functional Activities Questionnaire;^42^ autonomic symptoms using Compass-31;^43^ sleep disturbances using the Rapid Eye Movement Behaviour Disorder Sleep Questionnaire (RBDSQ)^44^ and visual hallucinations using the University of Miami PD hallucinations questionnaire (UMPDHQ). ^45^

Best corrected visual acuity, wearing habitual lenses if worn, was assessed using the LogMAR chart at 3m viewing distance. Contrast sensitivity was measured using the Pelli-Robson chart (SSV-281-PC) (http://www.sussex-vision.co.uk) at 1m viewing distance. Colour vision was tested using the Farnsworth D15 test^46^.

### 2.3 Plasma collection and processing

Approximately 30ml of blood was collected in polypropylene EDTA tubes. Samples were centrifuged at 2000g for 10 minutes, generating up to 16x0.5ml plasma and 14x0.5 ml serum aliquots, and stored immediately at -80°C.

NfL concentrations were measured using the Simoa Human Neurology 4-Plex A (N4PA) assay (Quanterix). P-tau217 concentration was measured using the AlzPath Simoa HD-X pTau-217 Advantage-PLUS kit^47^. All measurements were performed by analysts blinded to participant’ diagnoses and clinical data. P-tau217 measurements were performed in a single batch of reagents and NfL (4-Plex) measurement were performed in 4 batches of reagents.

### 2.4 MRI scanning

All MRI data was acquired using the same 3T Siemens Magnetom Prisma scanner using a 64 channel receive array coil (Siemens Healthcare, Erlangen, Germany). Participants underwent ∼1 hour of scanning, whilst receiving their usual medication. Structural anatomical scans consisted of T1 weighted MPRAGE images, acquired using 1×1×1□mm voxel, matrix dimensions 256×256×176, echo time (TE)=3.34ms, repetition time (TR)=2530ms, flip angle=7°. DWI was acquired using, 2×2×2□mm isotropic voxels, matrix dimensions 220x220x72, TE=3260ms, TR=58ms, b=50□s/mm^2^/17 directions, b=300□s/mm^2^/8 directions, b=1000□s/mm^2^/64 directions, b=2000□s/mm^2^/64 directions, acceleration factor=2.

### 2.5 Image analysis

#### 2.5.1 Preprocessing

Diffusion MRI images were pre-processed using Mrtrix3 by denoising,^48^ removal of Gibbs ringing artefacts,^49^ eddy current correction, motion correction^50^ and bias-field correction.^51^ Diffusion-weighted images were up-sampled to a spatial resolution of 1.3mm^3^.

#### 2.5.2 Fixel-based analysis of diffusion-weighted imaging data

Fibre orientation distributions (FODs) were computed using multi-shell 3-tissue-constrained spherical deconvolution using the group-average response function for each tissue type (grey matter, white matter & cerebrospinal fluid).^52,53^ Each participant’s FOD image was registered to a study specific template using affine and then nonlinear registration, and a template mask image calculated as the intersection of the brain masks for each participant.

Whole brain probabilistic tractography was performed on the FOD template to generate a tractogram with 20 million streamlines. To reduce tractography biases, this was then filtered to 2 million streamlines using spherical-deconvolution informed filtering of tractograms (SIFT).^54^

Fixel-based analysis was performed to characterise white matter tracts. For each participant, three measures were derived:

- Fibre cross-section (FC): a measure of macrostructure.
- Fibre density (FD): a measure of microstructural changes within tracts.
- Fibre density and cross section (FDC): a combined measure of change at both the micro-and macro-structural levels.^55^

Fixel-derived metrics were compared between people with DLB and PDD; LBD and PD-low risk, and LBD and controls, using age, sex and total intracranial volume as covariate (although not for comparisons involving FD, as recommended).^56,57^ We also performed correlational analyses across all people with LBD and PD, with motor and cognitive scores. All results are reported as *p* _FWE_<.05.

#### 2.5.3 Voxel-based analysis

A voxel-based analysis of fractional anisotropy (FA) and mean diffusivity (MD) was also performed. The diffusion tensor was calculated from the preprocessed diffusion images and used to derive FA and MD for each subject. The maps of each subject were registered to a study specific template and voxel-wise analysis conducted. Voxel-derived metrics was performed for the same comparisons and covariates as the fixel-derived metrics, using threshold-free cluster enhancement with default parameters^58^. All results are reported as *p* _FWE_<.05.

### 2.6 Other Statistical analyses

#### 2.6.1 Demographics & Plasma Markers

Demographic, cognitive and clinical measures were compared between groups using two-tailed Welch’s t-tests or Mann-Whitney tests for non-normally distributed data.

Plasma data were matched to clinical phenotype data, and a comparison made between disease groups, using one-way analysis of variance (ANOVA), controlling for age and sex. Planned comparisons were conducted to compare the combined LBD group with PD-low risk and control groups, and to compare the PDD versus DLB groups. Associations between plasma measures and cognitive/clinical scores were tested using linear regression, adjusted for age and sex. P<0.05, Bonferroni-corrected for multiple comparisons (p<.025 for group comparisons) was accepted as the threshold for statistical significance. Analyses were performed in R (R-4.2.1; https://www.r-project.org/).

## 3. Results

### 3.1 Demographics and clinical severity

A total of 47 DLB, 21 PDD, 43 PD, and 23 controls were recruited. 14 PD were classified as high-risk for developing dementia and were excluded, as they have been shown to have diffuse changes in white matter connections^8,15^. This left 29 PD-low risk.

The ages of people with DLB, PDD and controls did not differ significantly (mean ± std, DLB: 72.1 ± 5.8, PDD: 72.1 ± 6.4, control: 72.8 ± 6.8; DLB/PDD: t=0.20, p=.84; LBD/controls: t=0.585, p=.56), but PD-low risk (67.6 ± 5.1) were younger than people with DLB and PDD, t=3.59, p=.0006. There was a higher proportion of men with DLB and PDD than either PD-low risk (χ²=33.69, p<.0001) or controls (χ²=13.8, p=.0002). MoCA scores did not differ between people with DLB (21.3 ± 5.3) and PDD (21.8 ± 4.4), W=464.5, p=.80. As expected, MoCA scores were reduced for people with LBD compared to people with PD-low risk (28.9 ± 1.4; W=74, p<.0001) and controls (28.9 ± 1.1; W = 1492.5 p<.0001). Motor severity as measured using UPDRS-III was greater in people with LBD (PDD: 38.9 ± 1.11, DLB: 34.1 ± 17.6) than PD-low risk (26.0 ± 10.4; W=1338, p=.009) and controls (4.2 ± 3.5; W=20, p<.0001) groups. Demographics and clinical assessments are shown in Table 1.

**Table 1.** Demographic and clinical information of participants.

|  | PDD<br>(n = 21) | DLB<br>(n = 47) | Combined<br>LBD<br>(n = 68) | PD-low risk<br>(n = 29) | Controls (n<br>= 23) | PDD /<br>DLB | LBD / PD | LBD /<br>Control |
| --- | --- | --- | --- | --- | --- | --- | --- | --- |
| Age | 72.1 (6.43) | 72.1 (5.75) | 72.1 (5.91) | 67.6 (5.06) | 72.8 (6.78) | .85 | **<.0001** | .61 |
| Gender | 17 M / 4 F | 43 M / 4 F | 60 M / 8 F | 9 M / 20 F | 11 M / 12 F | .40 | **<.0001** | **.0001** |
| Years Education | 15.4 (3.58) | 15.0 (3.25) | 15.1 (3.33) | 16.2 (3.38) | 16.0 (2.31) | .73 | .17 | .32 |
| Disease specific metrics |  |  |  |  |  |  |  |  |
| UPDRS-III | 38.9 (11.1) | 34.1 (17.6) | 35.6 (15.9) | 26.0 (10.4) | 4.18 (3.51) | .14 | **.003** | **<.0001** |
| UPDRS-total | 80.2 (22.7) | 66.1 (30.1) | 70.4 (28.6) | 50.3 (18.9) | 6.48 (4.23) | **.024** | **.0008** | **<.0001** |
| LEDD | 705.2<br>(387.2) | 273.8(261.0<br>) | 406.6<br>(362.7) | 597.1<br>(343.2) | 0 (0) | **<.0001** | **.008** | **<.0001** |
| Disease Duration<br>Parkinsonism | 8.38 (5.57) | 2.15 (2.03) | 4.10 (4.55) | 7.04 (3.13) | - | **<.0001** | **<.0001** | - |
| Disease Duration<br>Dementia | 1.67 (1.98) | 2.17 (2.05) | 2.02 (2.03) | - | - | .022 | - | - |
| Global cognitive measures |  |  |  |  |  |  |  |  |
| MMSE | 26.5 (3.20) | 25.2 (3.88) | 25.6 (3.71) | 28.3 (5.52) | 29.2 (1.03) | .10 | **<.0001** | **<.0001** |
| MOCA | 21.8 (4.41) | 21.3 (5.34) | 21.5 (5.04) | 28.9 (1.37) | 28.9 (1.10) | .92 | **<.0001** | **<.0001** |
| Composite<br>Cognitive Score | -2.09 (1.57) | -2.77 (1.75) | -2.56 (1.71) | 0.20 (0.64) | 0.07 (0.47) | .15 | **<.0001** | **<.0001** |

All values are mean(standard deviation), apart from gender which is shown as a proportion. Combined LBD = Lewy body dementia (consisting of PDD and DLB. Note that these are the same patients as described in the separated DLB and PDD columns); DLB= Dementia with Lewy bodies; MMSE = Mini Mental State Examination, MoCA = Montreal Cognitive Assessment, PD-low risk= Parkinson's disease, at low risk of developing dementia; PDD = Parkinson's Dementia; UPDRS-III = Unified Parkinson's Disease Rating Scale part 3 (motor assessment). For people with PDD, disease duration is specified separately for onset of motor Parkinsonism and onset of dementia. Significant differences between groups are in bold.

**Table 2.** Cognitive and questionnaire scores.

|  | PDD<br>(n = 21) | DLB<br>(n = 47) | Combined<br>LBD<br>(n = 68) | PD-low risk<br>(n = 29) | Controls (n<br>= 23) | PDD /<br>DLB | LBD / PD | LBD /<br>Control |
| --- | --- | --- | --- | --- | --- | --- | --- | --- |
| **Cognitive measures** |  |  |  |  |  |  |  |  |
| Digit span backwards | 5.7 (2.0) | 5.5 (2.6) | 5.5 (2.4) | 8.2 (3.0) | 7.6 (2.3) | .59 | **.0001** | **.003** |
| (Half) Stroop colour<br>naming | 27.3 (9.2) | 28.2 (10.3) | 27.9 (10.0) | 17.5 (3.1) | 17.4 (2.5) | .62 | **<.0001** | **<.0001** |
| Stroop interference | 69.5 (48.5) | 71.1 (48.6) | 70.0 (48.1) | 31.5 (9.6) | 32.2 (5.4) | .78 | **<.0001** | **<.0001** |
| Verbal fluency category | 12.3 (4.8) | 10.3 (4.8) | 10.9 (4.8) | 22.3 (4.7) | 21.1 (5.4) | .11 | **<.0001** | **<.0001** |
| Graded naming task | 19.1 (5.5) | 21.0 (5.8) | 20.4 (5.7) | 25.1 (3.0) | 26.0 (3.2) | .12 | **<.0001** | **<.0001** |
| Verbal fluency letter | 13.0 (4.8) | 10.4 (5.4) | 11.2 (5.3) | 18.8 (4.7) | 17.7 (5.7) | .06 | **<.0001** | **<.0001** |
| Word recognition task | 21.6 (5.6) | 21.9 (3.0) | 21.8 (4.0) | 24.6 (0.9) | 24.6 (0.8) | .50 | **<.0001** | **<.0001** |
| Logical memory<br>(delayed) | 8.4 (4.3) | 7.0 (4.6) | 7.2 (4.5) | 15.1 (4.5) | 15.9 (4.0) | .36 | **<.0001** | **<.0001** |
| Hooper Visual<br>Organisation | 13.7 (7.1) | 17.7 (6.0) | 16.5 (6.5) | 26.2 (3.3) | 25.1 (2.4) | **.031** | **<.0001** | **<.0001** |
| **Clinical Questionnaires** |  |  |  |  |  |  |  |  |
| HADS Depression<br>Score | 6.8 (4.5) | 5.8 (3.2) | 6.1 (3.6) | 3.8 (2.8) | 2.8 (3.0) | .51 | **.003** | **<.0001** |
| HADS Anxiety Score | 8.2 (5.6) | 5.5 (3.9) | 6.2 (4.5) | 5.2 (3.4) | 4.4 (3.9) | .09 | .43 | .09 |
| Hallucinations | 2.2 (3.3) | 3.9 (3.0) | 3.4 (3.2) | 0.9 (2.5) | 0 (0) | **.019** | **<.0001** | **<.0001** |
| Sleep (RBDSQ) | 6.8 (3.9) | 7.4 (3.5) | 7.2 (3.6) | 4.1 (2.9) | 1.6 (1.5) | .45 | **.0002** | **<.0001** |
| Fluctuations (CAF) | 3.3 (3.2) | 5.9 (4.4) | 5.3 (4.3) | - | - | .09 | - | - |
| Functional activities | 11.6 (7.4) | 10.8 (6.6) | 11.0 (2.1) | 2.1 (3.0) | 0.2 (0.5) | .65 | **<.0001** | **<.0001** |

All values are mean(standard deviation). CAF=Clinician Assessment of Fluctuations; Combined LBD = Lewy body dementia (consisting of PDD and DLB. Note that these are the same patients as described in the separated DLB and PDD columns); DLB= Dementia with Lewy bodies; HADS=Hospital Anxiety and Depression Scale; PD-low risk= Parkinson's disease, at low risk of developing dementia; PDD = Parkinson's Dementia; RBDSQ=REM Sleep Behaviour Disorder Questionnaire. Significant differences between groups are in bold.

### 3.2 Plasma markers in LBD

#### 3.2.1 P-tau217

42 DLB, 20 PDD, 27 PD-low risk and 22 control participants had a sample available for inclusion in the plasma p-tau217 analysis. People with LBD showed higher concentrations of plasma p-tau217 than people with PD-low risk (*F*=10.80, *p*=.002) and age-matched controls (*F*=5.04, *p*=.028) (Figure 1A). P-tau217 concentrations did not differ between people with DLB and PDD (*F*=0.15, *p*=.70). Plasma p-tau217 was significantly associated with cognition, corrected for age and sex (MMSE: β= -4.62, p=.012; MOCA: β= -7.06, p=.0005; cognitive composite: β= -2.47, p=.002; Figure 1B and Supplementary Figure 2A and B), but was not associated with motor or total symptom scores (UPDRS-total: β=8.21, p=.75; UPDRS-III: β=1.65, p=.42).

**Figure 1.**
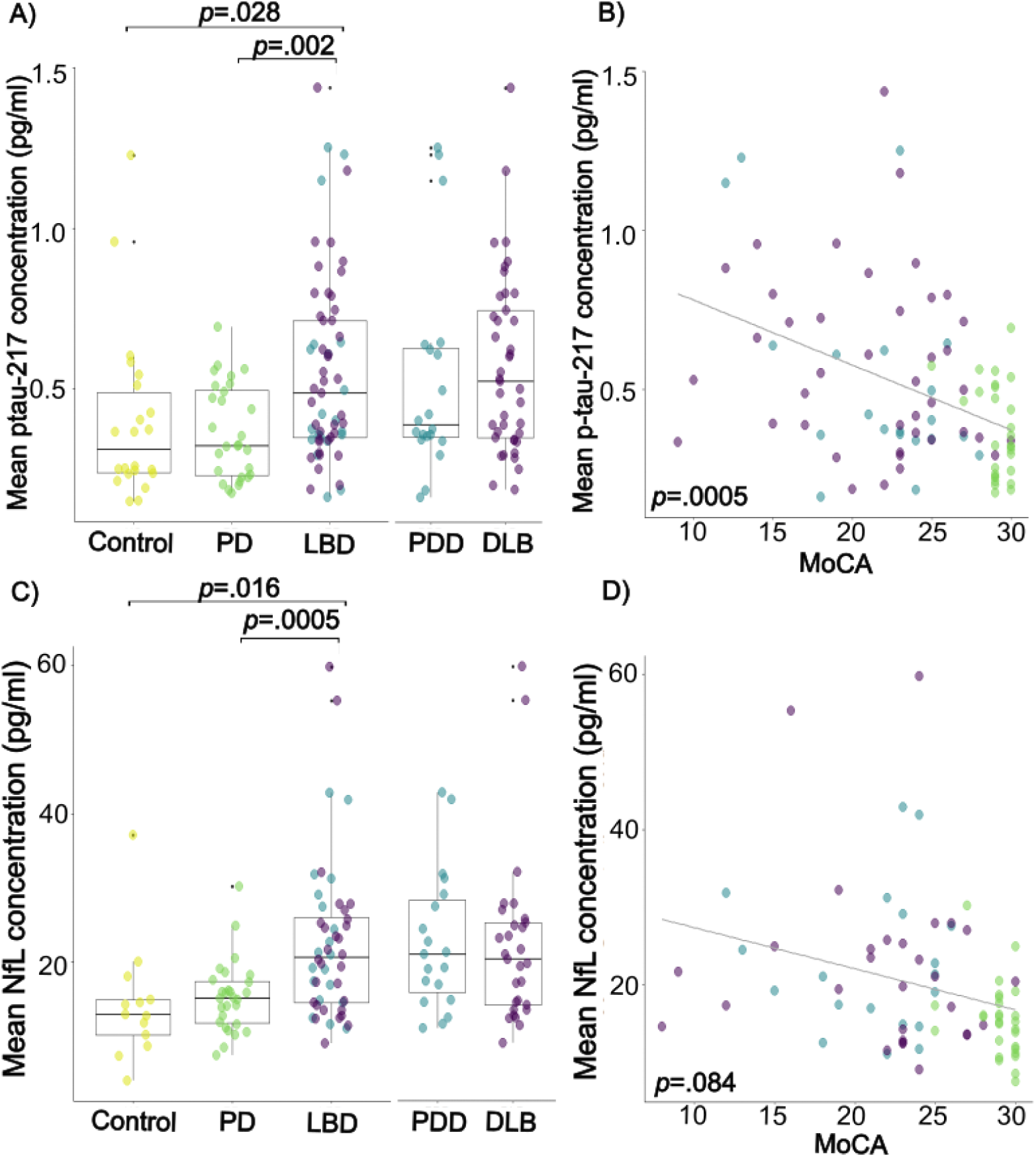
Plasma p-tau217 and neurofilament light (NfL) in Lewy body dementia (LBD). **A**) Differences in plasma p-tau217 concentrations between LBD, Parkinson’s low-risk and controls; and between LBD subtypes of Parkinson’s dementia (PDD) and dementia with Lewy bodies (DLB); **B**) Relationship between plasma p-tau217 and MoCA score in all patients (Purple = DLB, blue = PDD, green = PD-low-risk, as in panel 1A); **C**) Differences in plasma neurofilament light (NfL) concentration between LBD, Parkinson’s low-risk and controls; and between LBD subtypes of Parkinson’s dementia (PDD) and dementia with Lewy bodies (DLB); **D**) Relationship between NfL and MoCA score in all patients (Purple = DLB, blue = PDD, green = PD-low-risk, as in panel 1C). DLB=Dementia with Lewy Bodies, MoCA=Montreal Cognitive Assessment, PD=Parkinson’s Disease, LBD=Lewy Body Dementia, PDD=Parkinson’s Disease Dementia.

#### 3.2.2 NfL

30 people with DLB, 19 PDD, 27 PD-low risk and 13 control participants had a sample available for inclusion in the plasma NfL analysis. There was a significant effect of age on plasma NfL concentration (β=0.47, p=.011). There was no effect of reagent batch or sex. Mean concentration of plasma NfL was significantly greater in LBD than in PD-low risk (*F*=13.26, *p*=.0005, Figure 1C) and controls (*F*=6.36, *p*=.016), controlling for age and sex. Plasma NfL concentration did not significantly differ between people with DLB and PDD (*F*=2.56, *p*=.12). Plasma NfL concentration was significantly associated with the cognitive composite score (β= -0.06, p=.024, Supplementary Figure 1D), but not with any other clinical scores, corrected for age and sex (MMSE: β=-0.03, p=.66; MOCA: β=-0.11, p=.08; UPDRS-total: β=0.33, p=.35; UPDRS-III: β=0.15, p=.41; Figure 1D, Supplementary Figure 2C)

### 3.3 White matter connections in LBD

#### 3.3.1 Fixel-based analysis

##### 3.3.1.1 LBD vs PD-low risk

People with LBD showed reduced FC compared to PD-low risk in the genu of the corpus callosum, left anterior corona radiata and bilaterally in the sagittal stratum and posterior thalamic radiation (Figure 2A). No regions showed increased FC for LBD groups.

**Figure 2.**
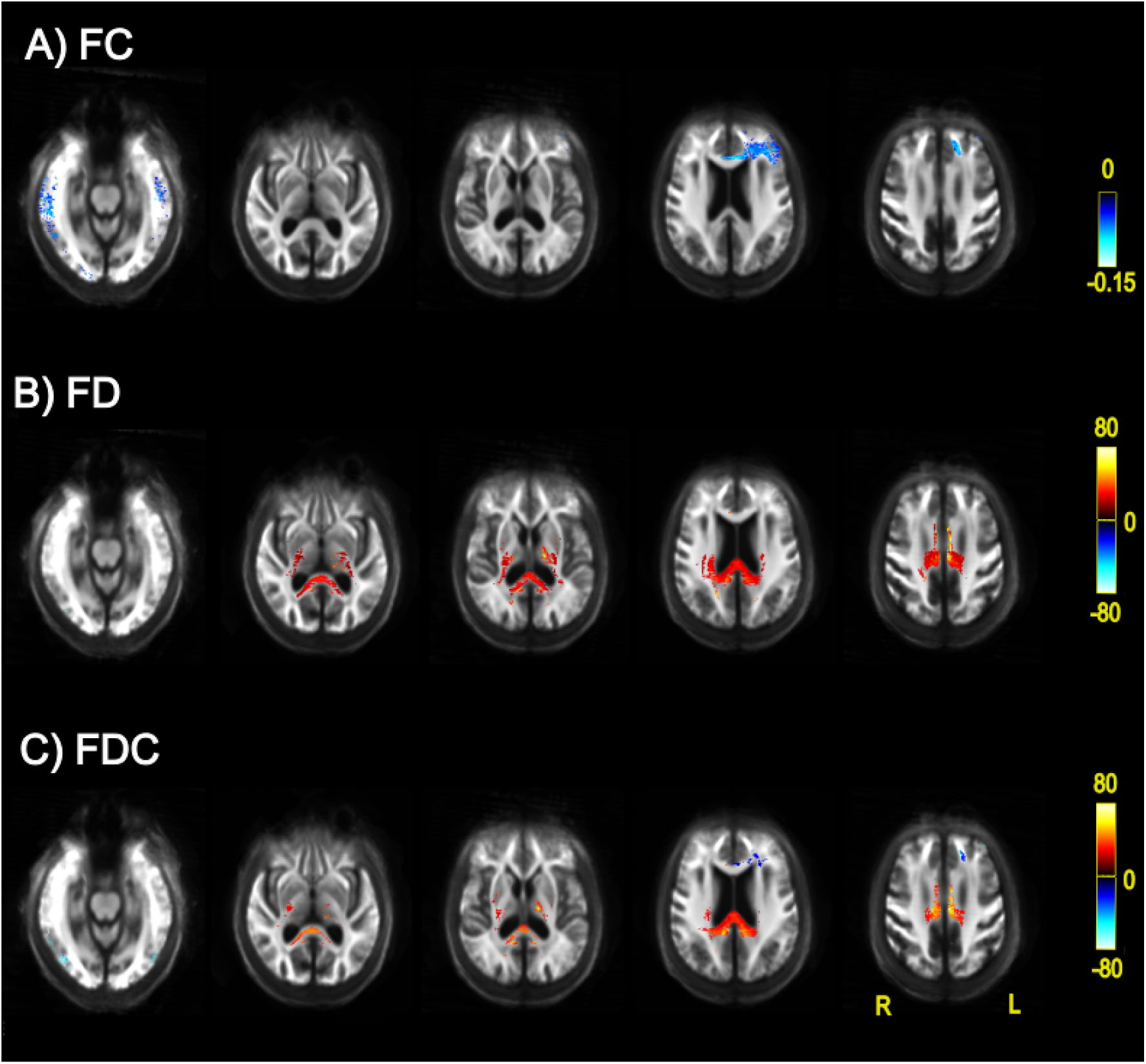
Differences in fixel-based metrics between people with Parkinson’s disease and Lewy body dementia. Percentage change for Lewy Body Dementia compared with Parkinson’s disease in **A**) Fibre density (FD), **B**) Fibre cross-section (FC), **C**) Combined fibre density and cross-section (FDC). Red-yellow colours indicate *increases* in fixel-based metrics and blue colours indicate *decreases* for LBD relative to the PD group. All results are displayed as streamlines corresponding to fixels that differed significantly between LBD and PD-low risk groups (*p* _FWE_<.05). Streamlines are displayed on the group white matter template and coloured by percentage change (colour bars).

FD was reduced for LBD relative to PD-low risk in temporal and occipital regions. FD was greater in the LBD group relative to PD-low risk in the genu, body and splenium of the corpus callosum, bilateral posterior limb and retrolenticular part of internal capsule, bilateral posterior and superior corona radiata, cingulum, right external capsule, right superior longitudinal fasciculus and bilateral cerebral peduncle (Figure 2B).

FDC was reduced for people with LBD relative to PD-low risk in the genu of the corpus callosum, bilaterally in the anterior corona radiata, and bilaterally in the sagittal stratum and posterior thalamic radiation. FDC was greater in the LBD group relative to PD-low risk regions including the body and splenium of the corpus callosum, bilateral posterior limb of internal capsule, bilateral posterior and superior corona radiata, cingulum, right superior longitudinal fasciculus and bilateral cerebral peduncle (Figure 2C).

##### 3.3.1.2 PDD vs DLB

People with PDD showed reduced FD compared to DLB in the body and splenium of the corpus callosum; cingulum; bilateral posterior limb and retrolenticular part of internal capsule; bilateral superior and posterior corona radiata; posterior thalamic radiation; left superior longitudinal fasciculus; left external capsule and left cerebral peduncle (Figure 3).

**Figure 3.**
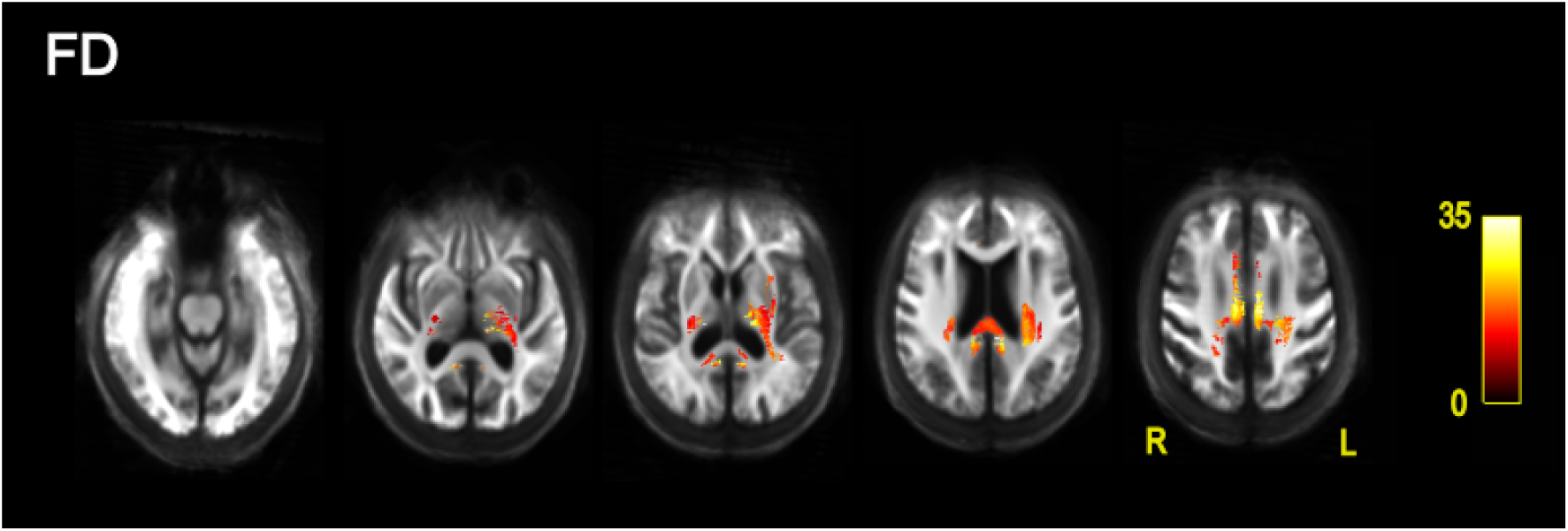
Fixel-based reductions in fibre density for Parkinson’s dementia relative to dementia with Lewy bodies. All results are displayed as streamlines corresponding to fixels that showed significantly reduced fibre density for the PDD relative to DLB group (*p* _FWE_<.05). Streamlines are displayed on the group white matter template and coloured by percentage change (colour bars).

There were no regions which showed reduced FD for people with DLB relative to PDD. No differences were observed in FC or in FDC between the two groups.

##### 3.3.1.3 LBD vs controls

No differences in fixel-based metrics were observed between LBD and control groups.

##### 3.3.1.4 Association with plasma concentrations and clinical variables

Higher concentrations of plasma p-tau217 were associated with reduced FC in people with LBD and PD in the splenium of the corpus callosum, fornix and bilaterally with the retrolenticular part of internal capsule (Figure 4), posterior thalamic radiation, sagittal stratum, and tapetum. Likewise, higher concentrations of plasma NfL were associated with reduced FC in the genu and splenium of the corpus callosum, posterior thalamic radiation, sagittal stratum and the right retrolenticular part of internal capsule. Neither plasma measure was associated with FD in people with LBD and PD.

**Figure 4.**
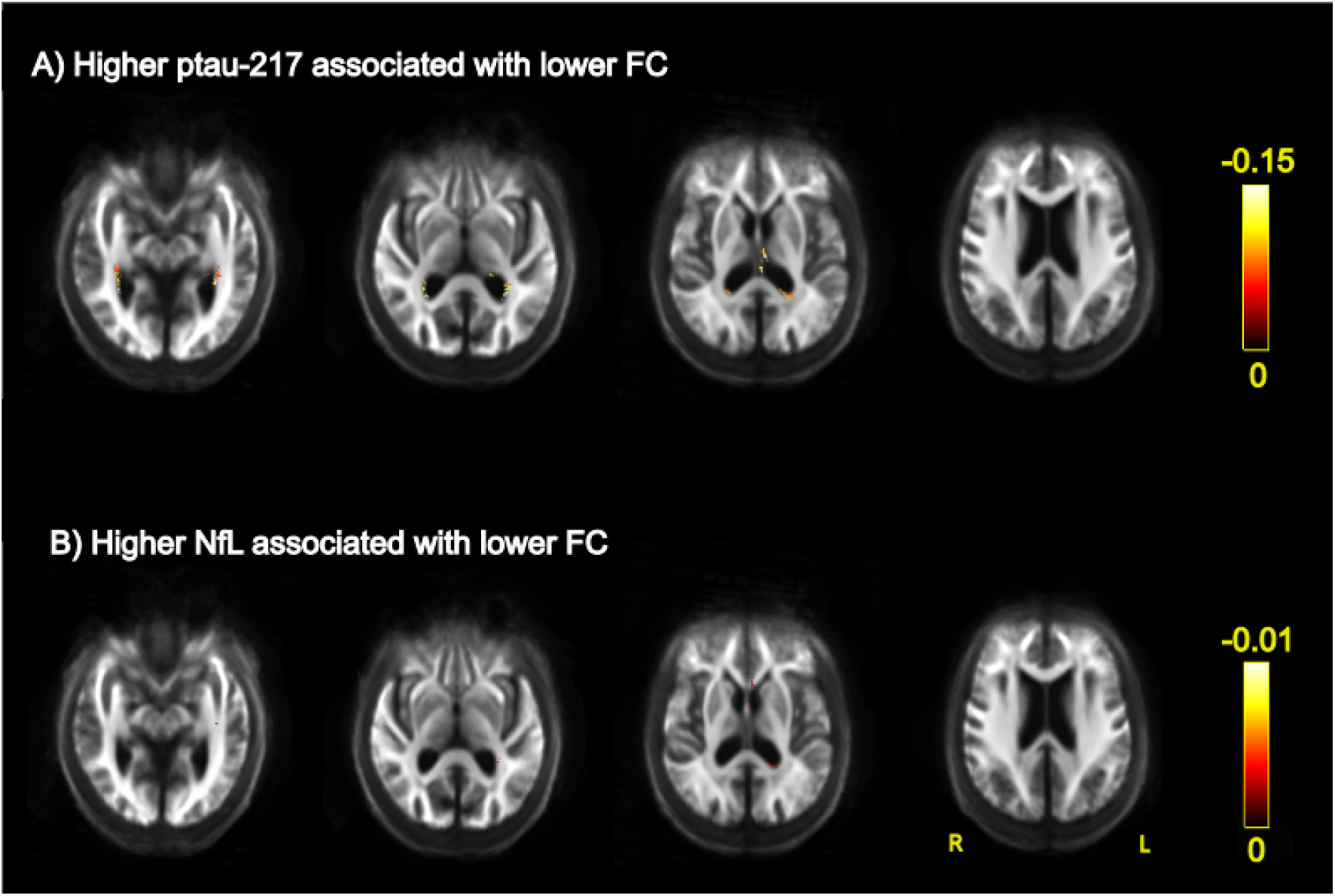
Associations between fixel-based fibre cross-section (FC) and plasma measurements. Association of **A**) increased concentrations of plasma p-tau217 with reduced fibre cross-section, **B**) increased concentrations of plasma neurofilament light (NfL) chain with reduced fibre cross-section. All results are displayed as streamlines corresponding to fixels that were significantly associated with plasma measures (*p* _FWE_<.05). Streamlines are displayed on the group white matter template and coloured by absolute effect (colour bars).

Significant associations were observed between FC and cognition in people with LBD and PD, although these were found in both directions (Figure 5): lower FC, indicating reduced white matter macrostructure, was associated with lower MMSE and MOCA bilaterally in the posterior thalamic radiation and with lower cognitive composite scores in the left sagittal striatum. Additionally, higher FD, indicative of preserved microstructure was associated with poorer MOCA score, but also with higher cognitive composite score in the left superior longitudinal fasciculus (Supplementary Figure 3). Higher UPDRS-III scores, reflecting worse motor symptoms were associated with lower FC in the cingulum (Supplementary figure 4). No association was seen between UPDRS-Total scores and FC or FD measures.

**Figure 5.**
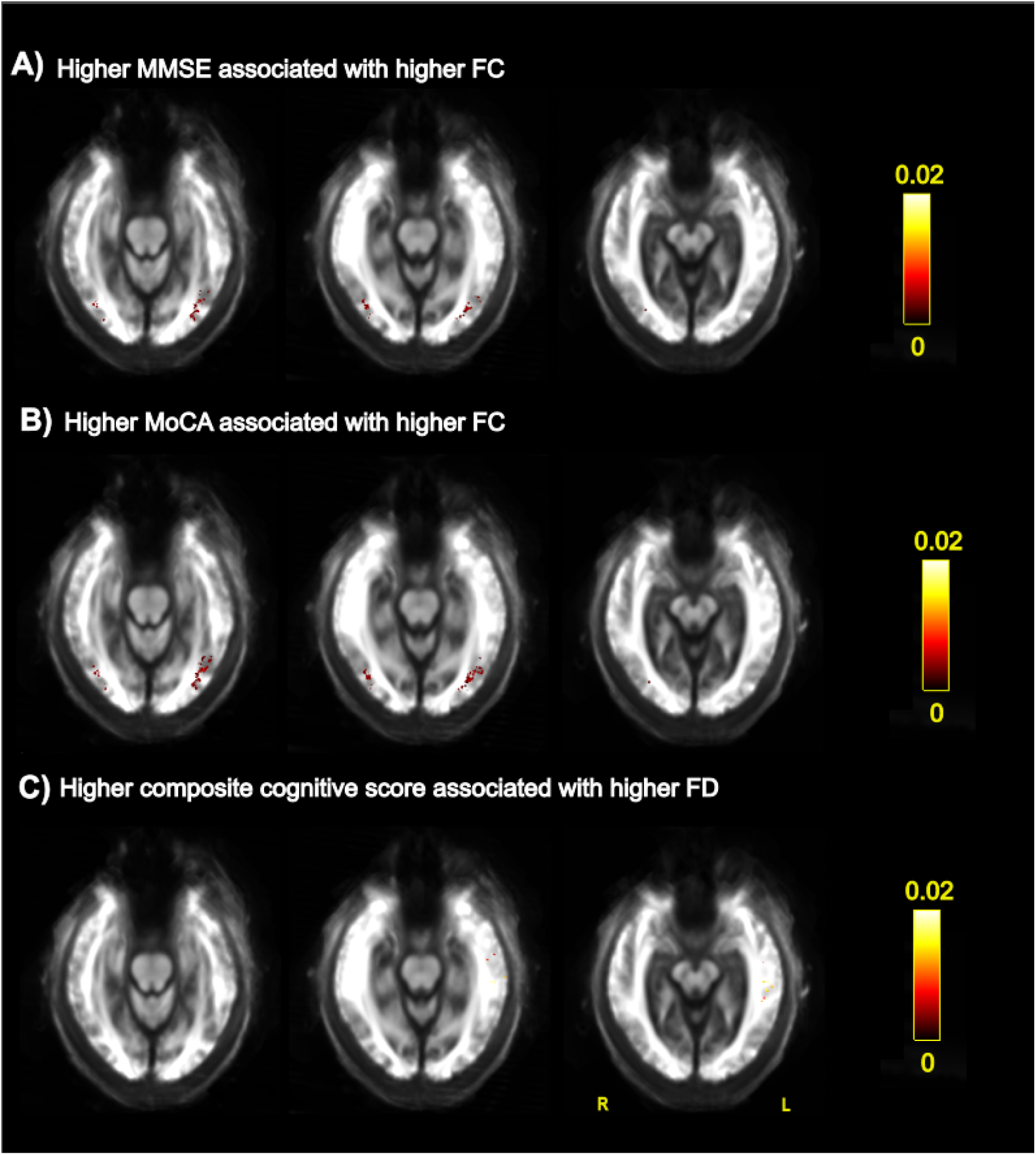
Associations between fixel-based fibre cross-section and cognitive measurements. Association of **A**) higher MMSE score with higher fibre cross-section, **B**) higher MoCA score with higher fibre cross-section, **C**) higher composite cognitive score with higher fibre cross-section. All results are displayed as streamlines corresponding to fixels that were significantly associated with cognitive measures (*p* _FWE_<.05). Streamlines are displayed on the group white matter template and coloured by absolute effect (colour bars).

#### 3.3.2 Voxel-based analysis

Conventional voxel-based analysis did not show any statistically significant differences between people with DLB and PDD after FWE correction. The combined LBD group showed increased mean diffusivity (MD) relative to people with PD-low risk, in widespread areas including the body and genu of corpus callosum, fornix, tapetum, superior fronto-occipital fasciculus bilaterally within the anterior, superior and posterior corona radiata, anterior and posterior limb of internal capsule, posterior thalamic radiation, superior longitudinal fasciculus, sagittal stratum and in the right cerebral peduncle (Figure 6). No differences in FA were observed between LBD and PD-low risk groups.

**Figure 6.**
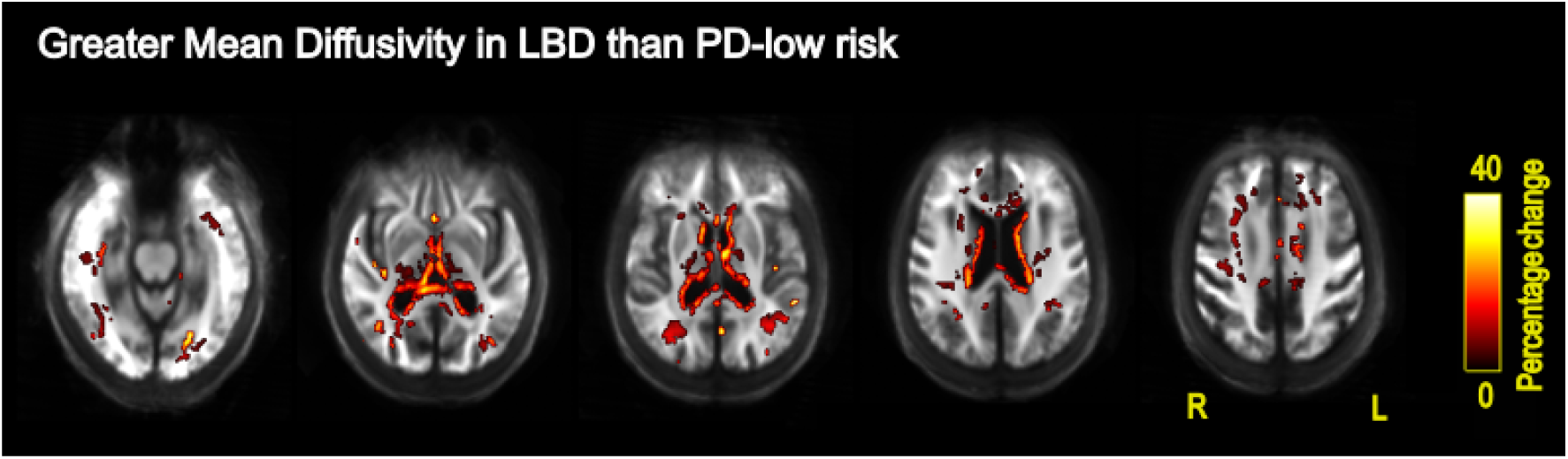
Voxel-based increases in mean diffusivity for Lewy body dementia relative to Parkinson’s disease with low dementia risk. All results are displayed as voxels that showed significantly increased mean diffusivity (MD) for the LBD relative to PD-Low risk group (*p* _FWE_<.05). Voxels are displayed on the group white matter template and coloured by percentage change (colour bars).

## 4. Discussion

We examined differences in diffusion imaging and plasma markers between people with LBD subtypes of DLB and PDD; and between people with LBD and people with PD who were low risk for dementia. We show macrostructural (fibre cross-section) changes, increased plasma NfL and increased plasma p-tau217 in people with LBD relative to PD-low-risk. We also found bidirectional microstructural changes (fibre density) for people with LBD relative to PD-low-risk. When comparing LBD subtypes, we found greater macrostructural changes (FC) in PDD relative to DLB, but no differences in plasma measures, or microstructural (FD) differences between LBD subtypes. This suggests that differences can be observed between people with DLB and PDD, using measures sensitive to white matter microstructure, and that these are distinct from differences between LBD and PD-low risk.

Our findings of reduced fibre cross section in LBD relative to PD-low-risk risk is consistent with our previous work: we used higher-order diffusion models and showed reduced fibre cross-section in PD-MCI and in PD at greater dementia risk ^15^, compared with people with PD who are low-risk for dementia. Other groups have used less-sensitive DTI approaches to compare LBD and PD and shown reduced FA in the corpus callosum, longitudinal fasciculus and fronto-occipital fasiculus for PDD and PD-MCI compared to PD with intact cognition.^10,11^ However, there are no other studies that apply whole-brain comparisons of the more sensitive higher order techniques to DWI. Similarly, DLB has not previously been investigated using fixel-based analysis.

Reductions in fibre-cross section are though to represent axonal loss, and reflect changes in area perpendicular to white matter bundles^14^. These changes have been linked with neurodegeneration and seem to be related to amyloid status^59^. Consistent with this observation, we found reductions in fibre cross-section for LBD compared with people with PD at low risk for dementia, where previous research shows greater neurodegeneration, measured using grey matter atrophy^60^ and higher rates of amyloid positivity than PD^61^. This is also supported in our work by higher plasma p-tau217 and NfL in LBD compared with PD-low-risk for dementia.

Our findings regarding plasma p-tau217 are noteworthy, as they represent, to our knowledge, the first report of plasma p-tau217 concentrations within LBD and PD. Our findings are consistent with previous research using earlier version of plasma p-tau (which are thought to be less sensitive). These showed increased p-tau181 relative to controls in DLB^22^ and PDD ^19^, as well as higher concentrations of p-tau181 associated with cognition in DLB^22^.

We also showed increased plasma NfL in LBD, consistent with previous work showing that NfL is increased in DLB^18^ and in PD with cognitive impairment^19,62^. NfL is a marker of axonal damage^17^, and we show that it is associated with changes in white matter macrostructure (FC). Interestingly, we did not find an association between NfL and MMSE or MoCA, which has been shown in some previous work ^19^ in LBD, but we did find an association of NfL with the composite cognitive score, which is more sensitive to subtle cognitive deficits.

Our finding of both increased and decreased fibre density for LBD relative to PD at low risk of dementia was unexpected, with increased fibre density in regions including the corpus callosum, internal and external capsule and decreases in temporal and occipital tracts. Some authors have suggested a role for compensatory white matter alterations in PD^63^, including in the cerebellum, internal and external capsule^64^. Another possible explanation relates to differences in small vessel disease (SVD) between PD and LBD groups. Fibre density is thought to reflect intra-axonal volume changes^14^. In a recent study of Alzheimer’s disease and genetic small vessel disease, microstructural differences were shown to be associated primarily with small vessel disease, with fibre density in Alzheimer’s strongly associated with white matter hyperintensities, lacunes and cerebral microbleeds across several tracts ^59^. SVD changes may differentially affect outcomes depending on their location. In the present study, reduced FD was seen in temporal and occipital regions. In these same regions, greater WMH burden has been shown to be associated with poorer cognitive function in older adults^65^, and with amyloid positivity in MCI ^66^. Likewise, in PD, enlarged perivascular spaces in temporal regions were shown to relate to cognitive scores, whilst those in the centrum semiovale did not ^67^. In our cohort, we did not have imaging sequences sensitive to small vessel disease available, but this could be examined in future work.

We found widespread changes in voxel-based mean diffusivity in people with LBD relative to PD patients at low risk of dementia, consistent with previous findings of changes in DTI metrics in both DLB^9,11,12^ and PDD ^10–12^. However, DTI models lack specificity, as they are unable to model crossing fibres and observed increases in mean diffusivity could be caused by multiple different processes, such as demyelination, axonal loss or inflammation^68^. Our fixel-based analysis instead allows us to interpret the observed MD changes with greater specificity and to show that both macro- and microstructural changes are relevant white matter differences observed between LBD and PD with low dementia risk. Indeed, we find bi-directional changes in fibre density in LBD versus PD which can not be appreciated using DTI-derived metrics. More detailed evaluation of metrics of higher specificity, such as fixel-based analysis, and how they relate with different pathological protein accumulation and other changes in tissue composition will be needed to better understand the drivers between white matter degeneration in LBD.

When we compared people with DLB and PDD, we found reductions in fibre density in people with PDD relative to DLB, but no differences were observed in DTI metrics. Whilst macrostructural differences were seen between LBD and PD groups, the difference in fibre density but not fibre cross-section metrics indicates predominantly microstructural differences between DLB and PDD. Fibre density differences were observed in regions including corpus callosum, corona radiata, internal capsule and superior longitudinal fasciculus. These are regions where white matter changes have previously been observed using DTI metrics in PDD and PD-MCI compared to PD and controls^10,11^. This suggests that in our cohort there were few differences between LBD subtypes that would be associated with amyloid status or neurodegeneration, but there may be differences in microstructure, associated with small vessel disease between people with DLB and PDD.

### 4.1 Limitations and Future directions

There are some potential limitations to consider within this study. We were unable to quantify concurrent small vessel disease, which would help to understand the changes in fibre density observed in our study. Future research should include fluid-attenuated inversion recovery or T2 MRI sequences to address this.

Our control and PD groups differed in age and sex to the LBD groups. Although we corrected for these factors in all analyses, it would have been preferable to recruit matched groups. All image acquisition was completed with participants taking their usual dopaminergic medication, in order not to affect cognitive scores. However, all comparisons are based on structural imaging measures, which are unlikely to be affected by dopaminergic agents.

### 4.2 Conclusion

In summary, we show reduced white matter macrostructure and increased plasma p-tau 217 and NfL in LBD compared with PD-low risk for dementia; and both of these measures were associated with poorer cognition. We also show poorer white matter microstructure for PDD relative to DLB. Our findings suggest both micro- and macro-structural changes in LBD, of which microstructure, implicating differences in small vessel disease may differ between LBD sub-types. Future work, sensitive to small vessel disease may help to further clarify these differences.

## Acknowledgements

We thank all the people with LBD, PD and controls who participated in this study. The authors acknowledge the use of the UCL Myriad High Performance Computing Facility (<u>Myriad@UCL</u>), and associated support services, in the completion of this work.

## Supplementary material

Supplementary material is available at [link to supplementary here].

