## Supplementary Methods for "Neuroimaging and plasma biomarker differences and commonalities in Lewy body dementia subtypes"

### Supplementary Materials

#### Supplementary Methods

If participants were unable to complete the full Stroop task, they completed a “Half-Stroop” consisting of the first 3 lines of the task. For participants who only completed a Half-Stroop, time to complete the full-Stroop colour naming was predicted using a regression model, derived from the full set of participant data, incorporated Half-Stroop time, age, sex and diagnosis. Missing full-Stroop times were predicted by combining the coefficients from this model with time to complete half-Stroop, age, sex and diagnosis. Predicted Stroop time was highly correlated with actual Stroop time in participants for which both measures were available (R^2^=0.97, Supplementary Figure 1), and predicted values were very close to actual values (RMSE = 4.65).

Additionally, if participants were unable to complete one task needed for the composite cognitive score, this was substituted with the mean z-score of that task for their disease group. If more than one task was not completed, the composite cognitive score was not calculated for that individual.

#### Supplementary Figures

**Supplementary Figure 1. Association between time taken to complete Stroop colour naming and predicted time based on time taken to complete half-Stroop colour naming.**

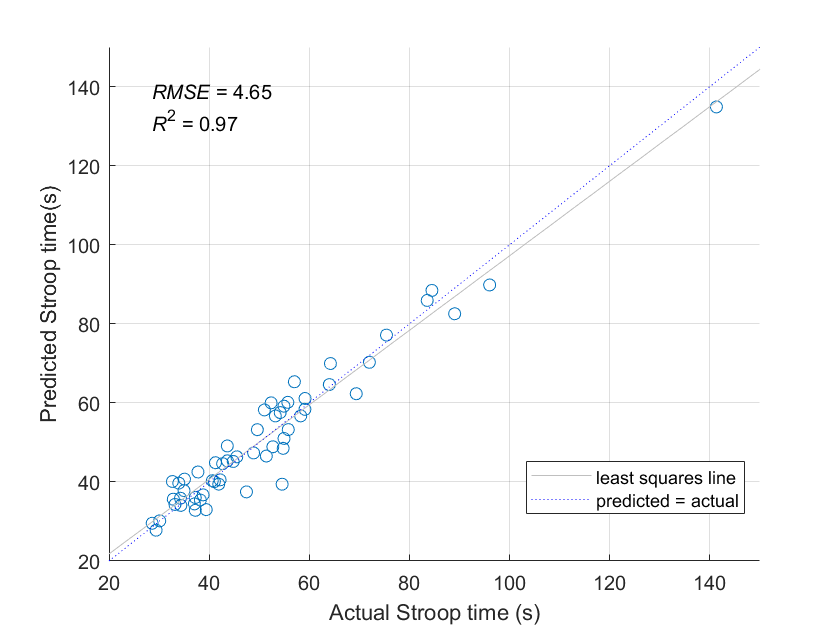

**Supplementary Figure 2. Plasma p-tau217 and NfL group associations with cognitive scores. A**) Association of plasma p-tau217 with MMSE score; **B**) Association of plasma p-tau217 with composite cognitive score. PD=Parkinson’s Disease, LBD=Lewy Body Dementia, PDD=Parkinson’s Disease Dementia, DLB=Dementia with Lewy Bodies, MMSE=Mini Mental-State Examination

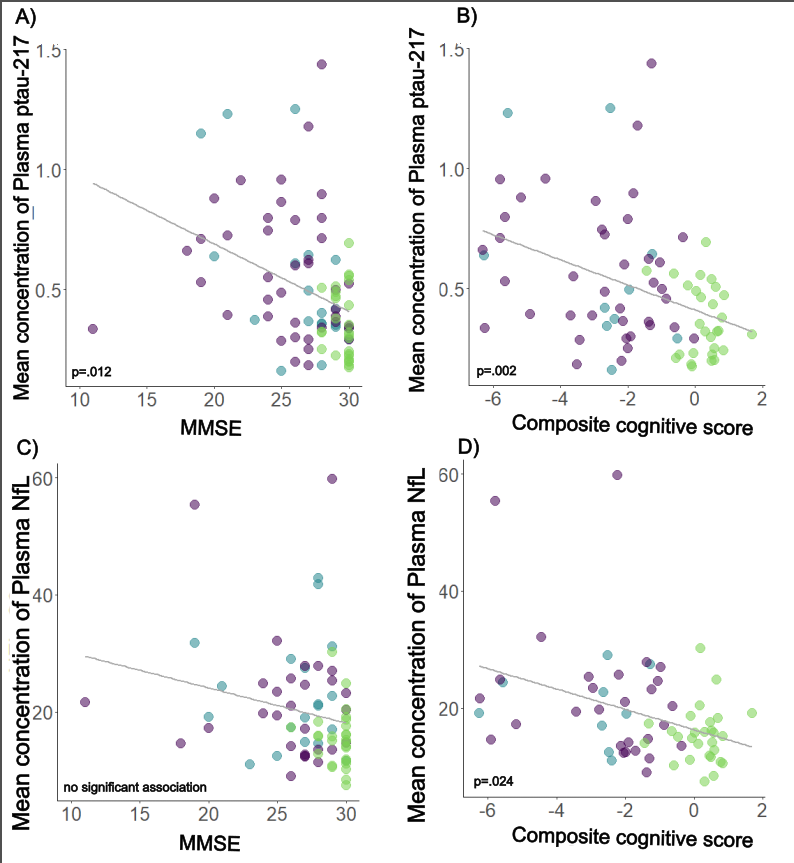

**Supplementary Figure 3. Associations between fixel-based fibre density and cognitive scores.** Association of **A**) higher MoCA score with reduced fibre density, **B**) higher MoCA score with higher fibre cross-section, **C**) higher composite cognitive score with higher fibre density. All results are displayed as streamlines corresponding to fixels that were significantly associated with cognitive measures (*p* _FWE_<.05). Streamlines are displayed on the group white matter template and coloured by absolute effect (colour bars).

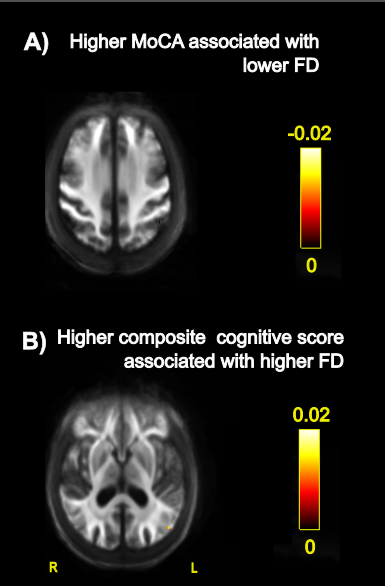

**Supplementary Figure 4. Associations between fixel-based fibre cross-section and UPDRS-III score.** All results are displayed as streamlines corresponding to fixels that were significantly associated with cognitive measures (*p* _FWE_<.05). Streamlines are displayed on the group white matter template and coloured by absolute effect (colour bars).

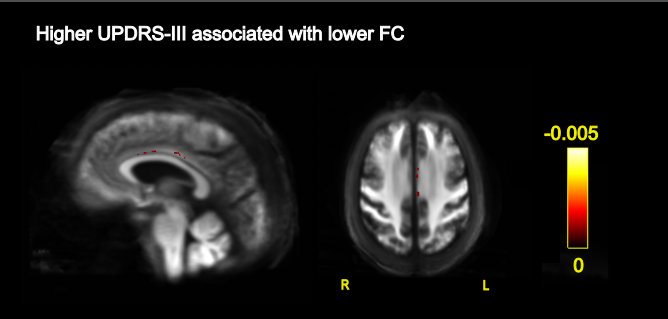
